# Dynamics of gut colonization by commensal and pathogenic bacteria that adhere to the intestinal epithelium

**DOI:** 10.1101/2024.10.24.620080

**Authors:** Dalaena E Rivera, Kayla Poirier, Samuel Moore, Ophélie Nicolle, Emily Morgan, Jonah-Faye Longares, Anupama Singh, Grégoire Michaux, Marie-Anne Félix, Robert J Luallen

## Abstract

Maintaining a healthy gut microbiome plays a critical role in avoiding gut-related pathologies. Bacterial adherence to the intestinal epithelium plays a vital role in niche establishment in the gut, as in vitro experiments and mathematical modeling suggest that adherence provides a strong competitive advantage over free-floating planktonic microbes. Currently, we lack the ability to study gut microbiome adherence in an in vivo model system. Through sampling natural populations of *Caenorhabditis*, we discovered three bacterial species that adhere to the intestinal epithelium of several wild *Caenorhabditis* isolates. When transferred to *C. elegans*, all three bacterial species colonized the entire anterior to posterior length of the intestine lumen. We isolated these bacterial species via in vitro growth or selective enrichment in the nematode gut and identified them as *Lelliottia jeotgali, Candidatus* Lumenectis limosiae, and *Candidatus* Enterosymbion pterelaium, the latter two representing new species. Adherent *Ca*. L. limosiae negatively affects host fitness, while *Lelliottia jeotgali* and *Ca*. E. pterelaium exhibited a neutral effect in our assays. We demonstrated that two of these species can actively proliferate in the intestine throughout the host lifespan, with *Lelliottia jeotgali* colonizing throughout the lumen simultaneously and *Candidatus* Lumenectis limosiae showing anterior-to-posterior directionality. In competition assays, animals pre-colonized with *L. jeotgali* significantly reduced colonization by pathogenic *Ca*. L. limosiae, but this effect was not seen when animals were colonized by both bacteria simultaneously. Strikingly, regardless of the colonization paradigm, populations exposed to both bacteria showed a near-identical mitigation of the pathogenic effects of *Ca*. L. limosiae. Altogether, these strains illustrate the capacity of microbiome bacteria to adhere, replicate, and establish a niche across the entire intestinal lumen in *C. elegans* and they present an opportunity to study bacterial adherence in the context of a whole, intact and transparent animal.

## Introduction

The intestinal epithelium plays a critical role in regulating the community of microbes in the gut. Dysregulation of this microbiome leads to gut dysbiosis, which is often implicated in a myriad of pathologies ranging from obesity to depression.^1,2^ Bacterial survival in the gastrointestinal tract is necessary to successfully establish a niche in the intestine, and epithelial cells play a critical role in providing selective pressures to control this community composition. These include negative pressures like host immunity and antimicrobial peptide production, and positive selective pressures, like secretion of metabolites favorable for beneficial bacterial species growth.^3,4,5,6.7^

While these pressures will affect bacterial survival in the gut lumen, niche formation and bacterial persistence are likely to be strongly affected by the ability to adhere to the intestinal epithelial surface.^8,9^ In fact, bacterial adherence to the mammalian intestinal epithelium is a common mechanism utilized by pathogens, like *Salmonella enterica* serovar Typhimurium and enterohemorrhagic *E. coli* (EHEC), and beneficial microbiome bacteria, like probiotic species of *Lactobacilli*.^10,11,12,13^ In vitro work suggests that more adhesive microbes can outcompete less adhesive genotypes, as bacteria with the capacity to adhere to microfluidics devices can displace non-adherent bacteria.^14^ Consequently, in these experiments planktonic, or non-adhering, bacteria are unable to colonize this occupied space.

According to mathematical modelling, even slow-growing adhering bacteria may have a competitive advantage over planktonic species due to better establishment of their niche within the intestine.^4^ However, we do not fully understand the role that adherence plays on governing bacterial colonization and niche establishment in the intestine. *Caenorhabditis elegans* has emerged as an excellent model to study host-microbe interactions in the gut.^15,16,17^ Aside from being genetically tractable and easy to maintain^18^, *C. elegans* shares numerous functional and morphological similarities to mammalian intestines. These include the presence of apical and basolateral polarization of intestinal cells, microvilli, and apical junctions that resemble mammalian cell-cell junctions.^19,20^ Yet, *C. elegans* intestines are simpler to study as they are comprised of a population of twenty non-renewable cells that are largely considered to be a single cell type.^20^ Due to *C. elegans* body transparency, intestinal colonization can be easily visualized in vivo using differential interference contrast (DIC) microscopy or RNA fluorescent in situ hybridization (FISH).^21^ Gut microbiome bacteria and intracellular pathogens, such as microsporidia and Orsay virus, naturally colonize and infect the intestines of wild-caught *C. elegans*.^17,22,23,24^ Recently, a simplified natural core microbiome was established in *C. elegans*, called CeMbio, providing the opportunity to study how the complexity of host-bacteria and bacteria-bacteria interactions influence microbiome composition and fitness.^22,25,26,27^ However, none of the twelve bacterial species comprising CeMbio are known to adhere to the intestine.^22^

Through ecological sampling of wild *Caenorhabditis* isolates, we discovered bacterial species that bind to the intestinal epithelial cells. These represent the first members of their gut microbiome capable of extensive intestinal adherence. Each of these bacterial species is horizontally transmitted amongst the host population and has the capacity to replicate and persist in the intestinal lumen without the need for constant bacterial uptake. We identified three distinct bacterial species in this collection, two members of Enterobacterales order and one member of Rickettsiales. We discovered that they have differential effects on host fitness, from neutral to negative, when colonizing independently. Co-colonization studies showed that adherence of a commensal bacterium prevented colonization of adherent pathogenic bacteria, though this had a detrimental effect on host reproductive fitness. Altogether, this unique set of bacteria presents an opportunity to utilize the *C. elegans* in vivo system to dissect the molecular mechanisms governing bacterial adherence to the gut, visualize the microbial biogeography in the intestines, and study the physiological impacts of bacterial adherence on the host.

## Results

### Discovery of adherent bacteria in the gut lumen of *Caenorhabditis* nematodes

By collecting wild nematodes in decomposing plant substrates, we find that they are associated with an array of microorganisms that likely form part of their natural microbiome.^22,25,26,27^ After sampling, single nematodes are picked onto standard NGM plates containing *E. coli* OP50-1 as food and their self-progeny are observed using differential interference contrast (DIC) microscopy. Over multiple years of sampling around the world, we have discovered multiple natural *Caenorhabditis* isolates that are colonized with microbes that adhere to the intestinal epithelium in the gut lumen (Table 1, Fig. 1A-1C). We categorized these adherent bacteria into three distinct morphological phenotypes: (1) bacilli that caused severe anterior distension of the host lumen, (2) thin bacilli that colonized the lumen with a hair-like appearance, and (3) bacilli that colonized with a club-shaped appearance. Fluorescent in situ hybridization (FISH) of these wild strains using a universal probe (EUB338)^21^ against the bacterial small ribosomal subunit (16S) rRNA revealed that all these microbes are bacteria (Fig. 1D-1F, left). After preliminary identification of these bacteria through 16S sequencing, we designed specific FISH probes to unique regions of each 16S and found that each of the above categories corresponded to distinct bacterial species or genera (Fig. 1D-F, right).

**Table 1.**
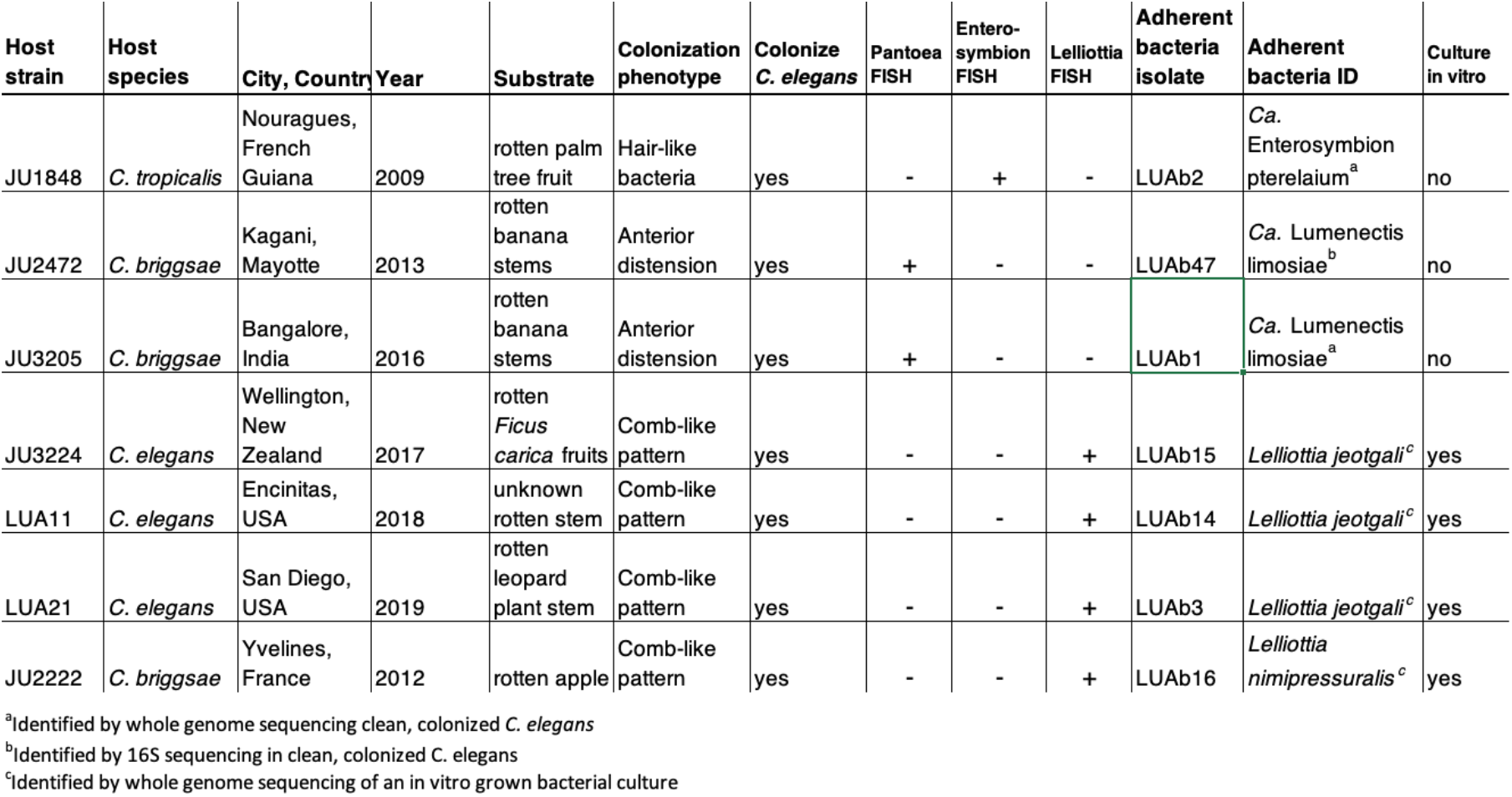
Collection of wild *Caenorhabditis* strains containing adherent bacteria.

**Figure 1.**
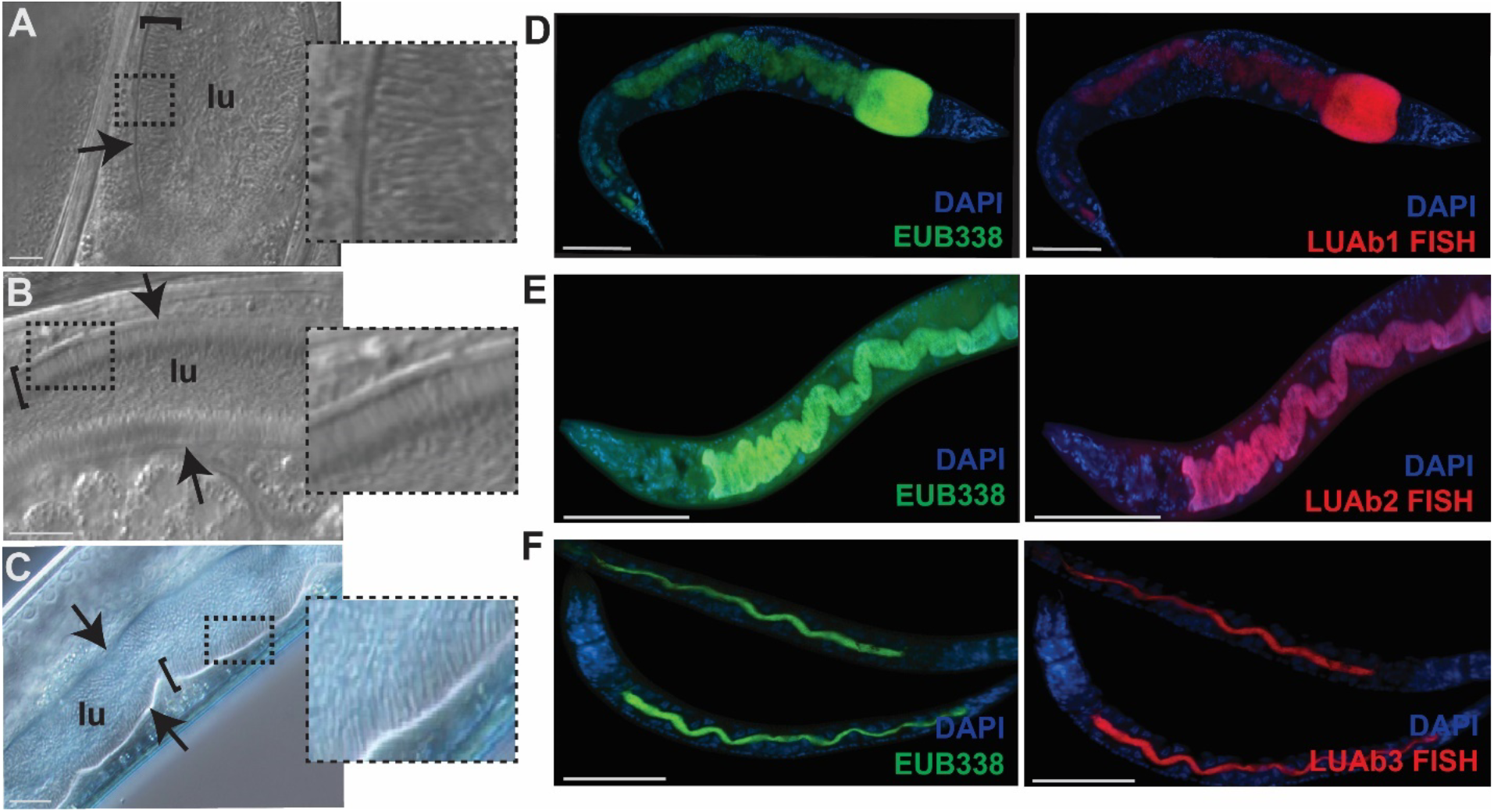
Discovery and visualization of three bacteria that adhere to the intestinal epithelium of wild *Caenorhabditis* isolates. (A-C) DIC microscopy images of the intestinal lumen (lu) of *Caenorhabditis* isolates demonstrating bacteria (indicated by brackets) adhering to the intestinal epithelium, denoted by arrows. Images to the right are insets of the dashed boxes seen on the left. (A) Adhering bacteria in *C. briggsae* (JU3205) that causes severe anterior distension of the lumen, later identified as *Ca*. Lumenectis limosiae (LUAb1). Scale bars are 10 μm. (B) Thin, hair-like bacteria in *C. tropicalis* (JU1848), later identified as *Ca*. Enterosymbion pterelaium (LUAb2). Scale bars are 5 μm. (C) Wild *C. elegans* (LUA21) colonized with comb-like adhering bacteria, later identified as *Lelliottia jeotgali* (LUAb3). Scale bars are 5 μm. (D-F) FISH using a universal 16S rRNA probe, EUB338 (green) and a species-specific FISH probe designed to the 16S rRNA (red) of either LUAb1, LUAb2, or LUAb3, respectively. Scale bars are 100 μm. Nuclei are stained via DAPI (blue).

We selected one representative wild strain of each morphological category and selectively cleaned it to enrich for the adherent bacteria.^28^ By forcing the animals into dauers, bacteria in the intestine are protected while external contaminants were removed through a harsh overnight wash of detergent and antibiotics. After enrichment, we assigned the designation LUAb1 to the bacteria that causes anterior distention, LUAb2 to the thin, hair-like bacteria, and LUAb3 to the club-like bacteria (Sup. Fig. 1A-C). LUAb3 was determined to be culturable in vitro after seeing persistent colonies on the NGM plates following selective cleaning. By contrast, the nematode strains containing LUAb1 (JU3205) and LUAb2 (JU1808) showed no obvious contamination on the plates after cleaning, suggesting that these bacteria were unculturable in vitro. Indeed, testing multiple culture media and techniques failed to result in bacterial growth for these two species.

To characterize these bacteria species, we conducted whole genome sequencing. LUAb3 was cultured in vitro while LUAb1 and LUAb2 were grown in the host gut lumen (*C. elegans* N2). After de novo assembly of their genomes and phylogenomics, we newly describe LUAb1 as a novel bacterium in the Enterobacterales order we named *Candidatus* Lumenectis limosiae, due to the characteristic distension in colonized animals (Fig. 2A, Sup. Text 1, Taxonomic Summary). We newly describe LUAb2 as a novel bacterium belonging to the Rickettsiales order that we named *Candidatus* Enterosymbion pterelaium after the King Pterelaus in Greek mythology (Fig. 2B, Sup. Text 1, Taxonomic Summary). Finally, we identified LUAb3 as an isolate of the existing species *Lelliottia jeotgali* in the *Enterobacteriaceae* family (Fig. 2A).^29^

**Figure 2.**
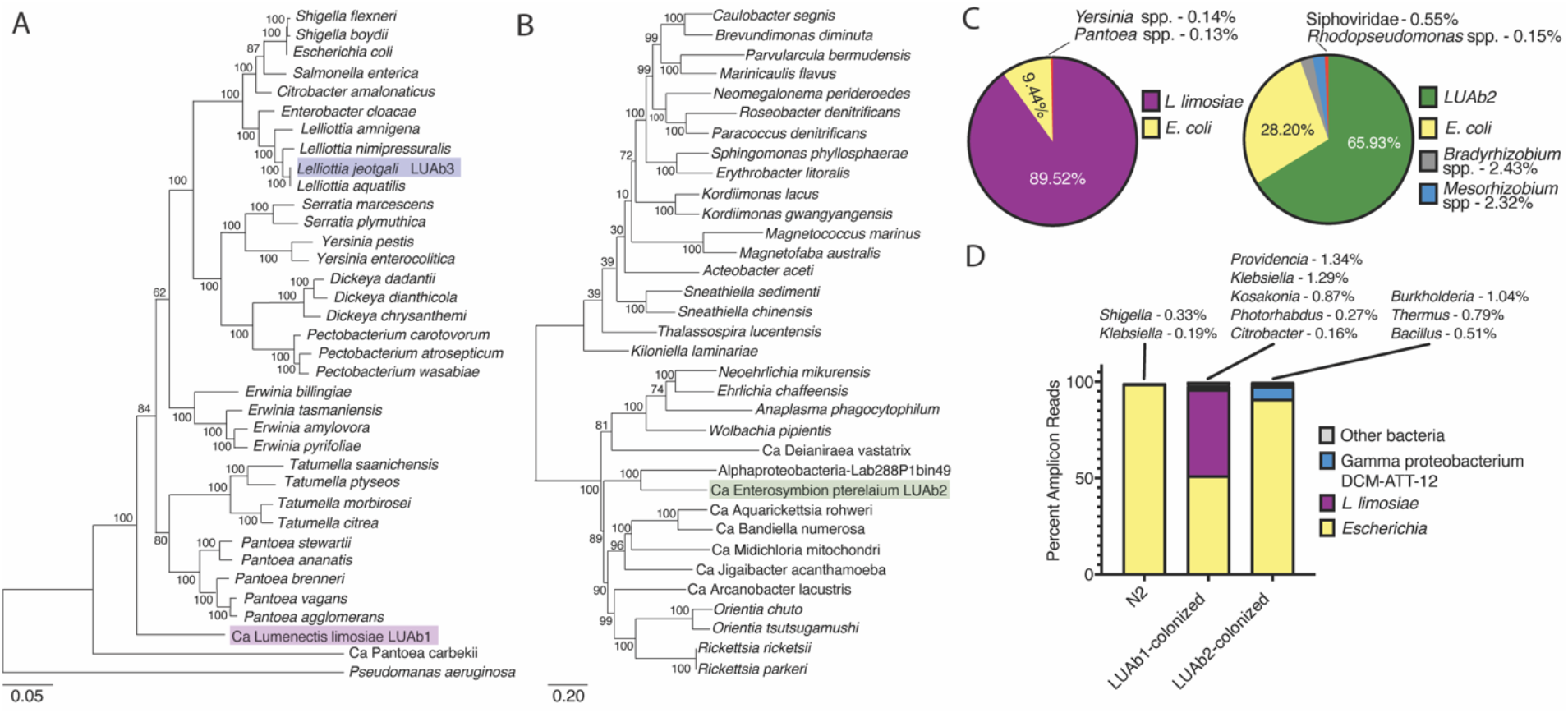
Phylogenomic identification of adherent bacteria species. (A) Phylogenomic tree of sequenced Enterobacterales spp. with outgroup *Pseudomonas aeruginosa*. Branch lengths are the number of substitutions per site and branch points indicate percentage of trees with clustering of associated taxa. (B) Phylogenomic tree of sequenced Alphaproteobacteria spp. Branch lengths are the number of substitutions per site and branch points indicate percentage of trees with clustering of associated taxa. (C) Sequenced whole genomic DNA from clean *C. elegans* ERT413 colonized with LUAb1 or LUAb2 were analyzed via metagenomics. Pie charts show the percent of non-*C. elegans* reads that were identified to belong to a particular bacterial species or genus. (D) 16S PCR and amplicon sequencing of *C. elegans* ERT413 colonized with LUAb1 or LUAb2, with uncolonized N2 as a control. Pie charts show the percent of non-amplicon reads that were identified to belong to a particular bacterial species or genus.

Members of Rickettsiales are typically obligate intracellular parasites.^30^ *Ca*. E. pterelaium does not show any evidence of intracellular invasion (0% invasion incidents out of 101 animals across 3 independent replicates, scored in fully colonized animals stained by FISH). However, we have found that this bacterium does not survive well when separated from the host through host lysis and filtering (see below). Its membership in Rickettsiales may explain the obligate-like nature of this bacterium. *Ca*. L. limosiae and *L. jeotgali* fall within the Enterobacterales order, which contains both commensals and pathogens, with the most notable species being *Escherichia coli*.^31^ Amongst the four adhering *Lelliottia* isolates, LUAb3, LUAb14, and LUAb15 are *L. jeotgali* and LUAb16 is *L. nimipressuralis* (Table 1). *Lelliottia* species are often associated with water sources, food, plants, and in some cases human clinical samples.^32-35^ Both *L. nimipressuralis* and *L. amnigena* are known plant pathogens causing soft rot in dangshen roots and onion bulb decay, respectively.^34,35^ This potentially explains how wild *Caenorhabditis* nematodes may encounter these *Lelliottia* species. Interestingly, in the same genus, *Lelliottia amnigena* JUb66 is a member of CeMbio^25^ and was also found colonizing the *C. elegans* cuticle^36^ but does not adhere to the gut.

Despite not being able to culture *Ca*. L. limosiae or *Ca*. E. pterelaium in vitro, several results indicate that we were able to remove a majority of contaminating bacteria and establish a population of *C. elegans* N2 that remains persistently colonized. First, after cleaning as described above, the cultures of *C. elegans* N2 reference strain containing either *Ca*. L. limosiae or *Ca*. E. pterelaium showed no obvious external contamination growing on the NGM plates. Second, when comparing species-specific FISH to generic FISH (EUB338), we saw no detectable colonization in the gut by any other species (see Fig. 1D-E). Third, we analyzed the read distribution after whole genome sequencing in *C. elegans*. For *Ca*. L. limosiae, we found that the vast majority of non-*C. elegans* reads were either from *E. coli* (OP50-1) or *Ca*. L. limosiae, with ∼1% of reads distributed to other microbes. Similar results were seen for *Ca*. E. pterelaium, with ∼6% of reads coming from other microbes (Fig. 2C). Fourth, similar levels of microbial contamination were seen when we conducted 16S amplicon sequencing in clean *Ca*. L. limosiae- and *Ca*. E. pterelaium-colonized strains. *E. coli* reads were predominant in the *Ca*. E. pterelaium-*C. elegans* culture likely due to the 16S PCR primers used that target Gammaproteobacteria and do not amplify Alphaproteobacteria efficiently (Fig. 2D). These data show that despite not being able to culture *Ca*. L. limosiae or *Ca*. E. pterelaium in vitro, we were able to remove a majority of contaminating bacteria and establish a population of *C. elegans* N2 that remains persistently colonized with these microbiome bacteria.

Finally, we tested our collection of wild *Caenorhabditis* strains with adhering bacteria using *Ca*. L. limosiae, *Ca*. E. pterelaium, and *L. jeotgali* specific FISH probes to determine their relative relationship to these bacteria. We found that four nematode isolates from France, New Zealand, and the USA were colonized with *Lelliottia* spp. similar to *L. jeotgali* LUAb3, two from India and Mayotte were colonized with a *Ca*. Lumenectis spp., and only one isolate from French Guiana was colonized with a *Ca*. E. pterelaium-like bacteria (Table 1). Overall, we have isolated and identified three divergent adherent bacterial species found in the intestinal lumen of wild-isolated *Caenorhabditis* nematodes from around the globe.

### Contrasting effects of three adherent bacteria on host reproductive fitness and physiology

To determine whether these adhering bacteria affect *C. elegans* fitness, we conducted lifespan and brood size assays. To better visualize the colonization within the intestine, we used *C. elegans* strain ERT413 expressing GFP specifically in intestinal cells^37,38^ We assayed the effects of the bacteria on the host reproductive fitness, specifically on reproductive lifespan and brood size. We found that ERT413 animals colonized with LUAb1 (*Ca*. L. limosiae) exhibited a substantial decrease in reproductive lifespan compared to uncolonized ERT413, with only ∼20% of animals surviving to day 8 of adulthood compared to ∼80% of non-colonized controls (Fig. 3A). By contrast, animals colonized with LUAb2 (*Ca*. E. pterelaium) or LUAb3 (*L. jeotgali*) do not appear to affect host lifespan compared to controls (Fig. 3B-C). A similar result was observed with brood size assays, as LUAb1 colonized animals showed a ∼45% decrease in total brood size (Fig. 3D), while LUAb2 and LUAb3 did not show a significant decrease (Fig. 3E-F). There was a small, significant decrease in brood for LUAb3-colonized animals when specifically looking at daily brood sizes, but this was at a time point after most of the brood had been laid (day 3 and day 4, Fig. 3F). Overall, this data suggests that under these conditions, *Ca*. L. limosiae is a pathogen to *C. elegans*, while *Ca*. E. pterelaium and *L. jeotgali* behave as commensal-like microorganisms.

**Figure 3.**
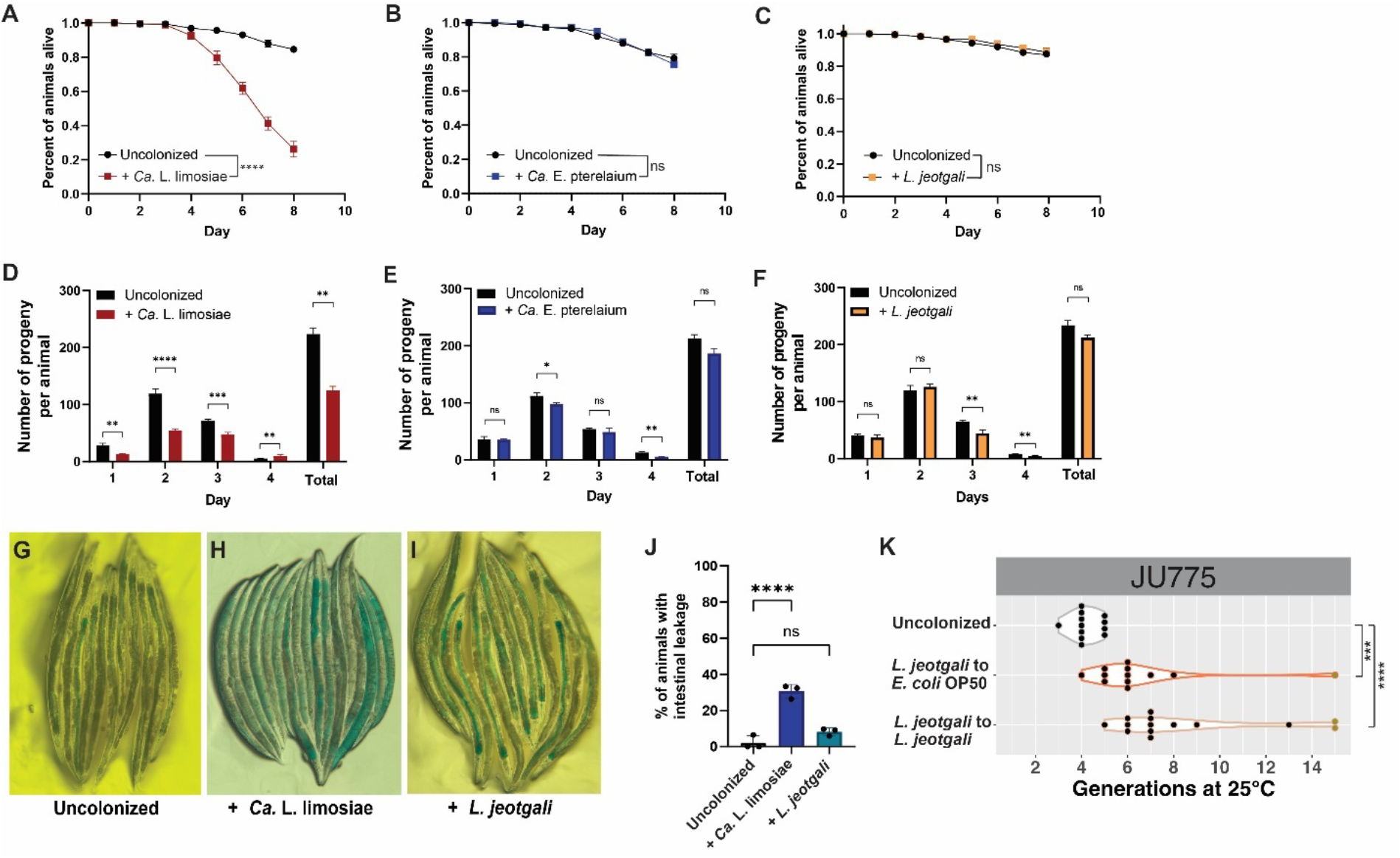
Adhering bacteria have differing effects on *C. elegans* fitness. (A-C) Reproductive lifespan of ERT413 colonized with *Ca*. L. limosiae, *Ca*. E. pterelaium, or *L. jeotgali*, respectively. n=60 in 3 independent experiments. Error bars show standard deviation (SD) from the mean. Statistical analysis performed by Mantel-Cox test, where p < 0.0001 = **** and ns= not significant. (D-F) Brood size of ERT413 colonized with *Ca*. L. limosiae, *Ca*. E. pterelaium, or *L. jeotgali*, respectively, n=60 in 3 independent experiments. Statistical analyses performed by unpaired two-tailed t-test where ns = not significant, p < 0.05 = *, p < 0.01 = **, p < 0.001 = ***, and p < 0.0001 = ****. (G-I) Representative images for “Smurf” assay testing of intestinal barrier function assay. (G) Uncolonized *C. elegans* N2 fed E. coli OP50. (H) *C. elegans* N2 colonized with *Ca*. L. limosiae. (I) *C. elegans* colonized with *L. jeotgali*. (J) Quantification of G-I, showing percent of animals that experienced leakage of blue dye from the intestine into the germline and/or body cavity, where N=3 and n=87-108. Statistical analyses performed by ordinary one-way ANOVA where ns = not significant and p < 0.0001 = ****. (K) Mortal germline (Mrt) phenotype describing the number of generations until sterility of wild *C. elegans* strain JU775 either fed only *E. coli* OP50 (grey), colonized first with *L. jeotgali* LUAb3 and transferred to *E. coli* OP50 (orange), or fed only *L. jeotgali* LUAb3 (brown). Wilcoxon rank tests p<0.001 = *** and p<0.0001 = **** across at least 12 replicates for each of the three conditions.

Next, we measured the intestinal barrier function of *C. elegans* when colonized with *Ca*. L. limosiae and *L. jeotgali* through a “Smurf” assay in which animals are fed a blue dye that can pass into the body cavity if the intestinal epithelia is compromised .^39,40^ Due to difficulties culturing *Ca*. L. limosiae in vitro, we adopted LUAb1 (*Ca*. L. limosiae) for an existing protocol used with microsporidia spores to better control the amount of exposure to the bacteria.^24^ Unfortunately, we were unable to prepare LUAb2 (*Ca*. E. pterelaium) in a similar manner. Overall, we found that at 2 days post colonization *C. elegans* N2 adults colonized with *Ca*. L. limosiae exhibited an increase in intestinal leakiness resulting in the presence of blue dye in the body cavity compared to uncolonized controls fed only OP50 (Fig. 3G-H). On the other hand, there was no significant difference in intestinal leakage between *L. jeotgali* colonized animals and the uncolonized control (Fig. 3I-J).

Finally, we tested if *L. jeotgali* (LUAb3) could suppress the mortal germline (Mrt), a phenotype seen in some *Caenorhabditis elegans* wild isolates where the lineage becomes sterile after several generations at 25°C. Previously, another *L. jeotgali* isolate (JUb276) was demonstrated to suppress the Mrt phenotype^41^ and Orsay virus infection.^42^ We found that colonization with *L. jeotgali* LUAb3 could also suppress the Mrt phenotype of wild *C. elegans* strain JU775 when grown at high temperatures, demonstrating a beneficial effect (Fig. 3K). Altogether, these data further support the pathogenic nature of *Ca*. L. limosiae and the commensal or beneficial properties of *L. jeotgali* on *C. elegans*.

### Adhering bacteria cause differing effects on physiology and structure of the host intestine

Because of the proximity between the host intestinal epithelial cells and the adhering bacteria, we aimed to better visualize this interaction and its physiological impacts on the host through transmission electron microscopy (TEM), focusing on *Ca*. L. limosiae- and *L. jeotgali*-colonized *C. elegans*. In N2 uncolonized animals fed OP50 bacteria, we did not see live bacteria in the gut lumen, but rather circular, electron-dense structures of various sizes that may represent digested bacteria (Fig. 4A). The intestinal cells contain characteristically long microvilli surrounded by a moderately electron-dense glycocalyx. By contrast, animals colonized with *Ca*. L. limosiae (LUAb1) contained intact bacteria, with a clustering of bacterial cells directly adjacent to the glycocalyx of dramatically shortened intestinal microvilli (Fig. 4B). Intact bacterial cells were also observed in the lumen of animals colonized with *L. jeotgali* (LUAb3), some of which were also directly adjacent to the intestinal glycocalyx (Fig. 4C). We did not observe the organized comb-like adherence pattern seen via DIC microscopy (Fig. 1C), although this may be attributed to the orientation of the focal plane as dark bacterial structures can be seen along the perimeter of the intestinal epithelia near the glycocalyx.

**Figure 4.**
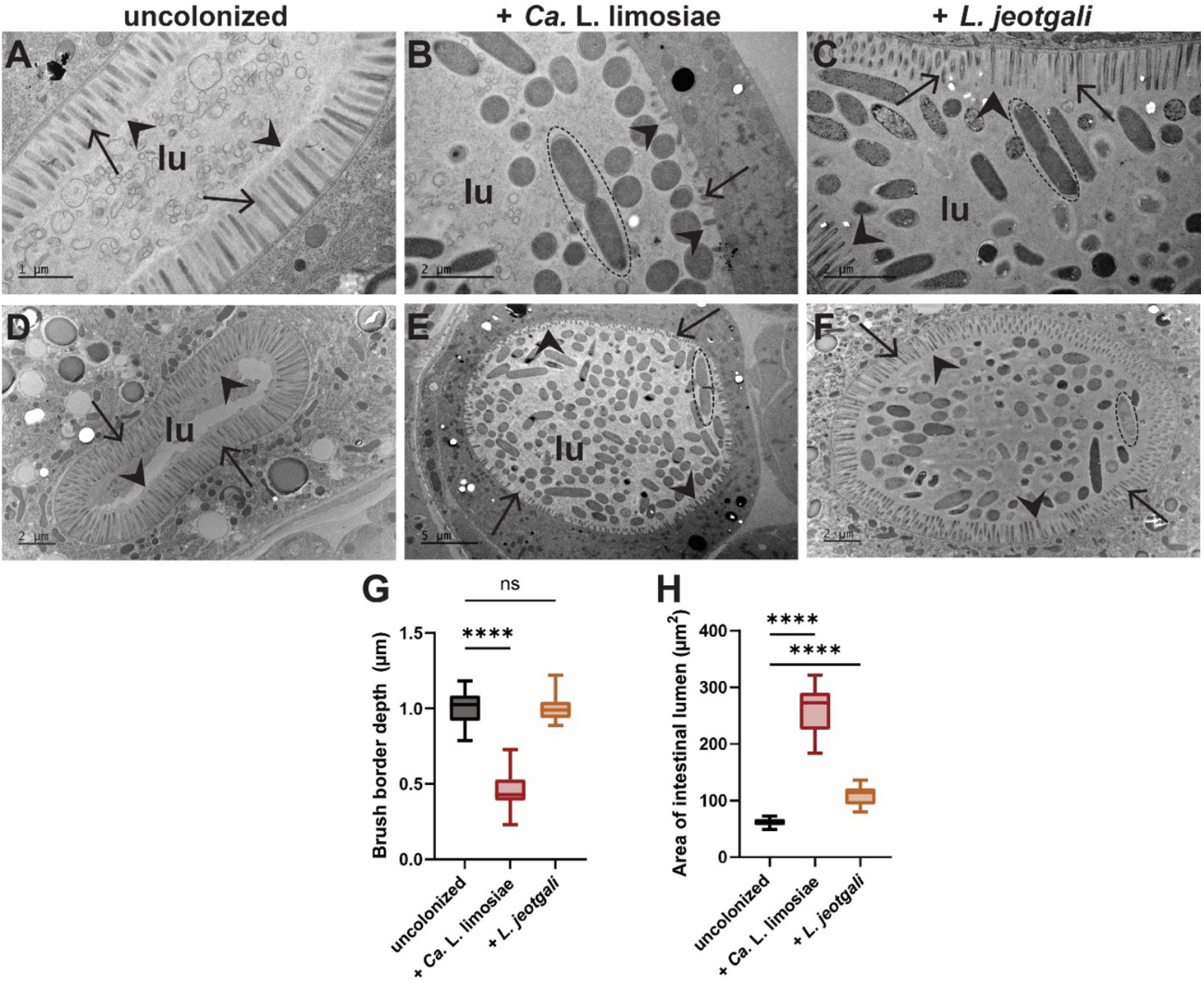
Visualization of adhering bacteria in the host intestine by TEM. lu indicates the intestinal <u>lu</u>men, arrowheads indicate the apical side of the glycocalyx, and arrows denote the microvilli. Bacteria undergoing septation are indicated by a dashed oval. (A-C) Representative TEM images of *C. elegans* intestine colonized with adhering bacteria (A) Intestine of uncolonized *C. elegans* N2. Scale bars are 1 μm. (B) *C. elegans* ERT413 (*jySi21[spp-5p::GFP; Cbr-unc-119(+)] II*) colonized with *Ca*. L. limosiae seen as the dark rod and spherical structures in the lumen, scale bars are 2 μm. (C) Image of the intestine of *L. jeotgali* colonized *C. elegans* N2, with scale bars = 1 μm. Individual *L. jeotgali* cells are seen as dark round or ovoid structures in the lumen. Dividing bacteria are outlined by a dashed oval. Scale bars indicated are 1 μm. (D-F) TEM images showing the intestinal distension in *C. elegans* N2 colonized with *Ca*. L. limosiae and *L. jeotgali* in the intestine, respectively. (D) Cross section of intestine of uncolonized *C. elegans* N2. Scale bars are 2 μm. (E) *C. elegans* ERT413 (*jySi21[spp-5p::GFP; cb-unc-119(+)] II*). Scale bars are 5 μm. (F) Image of the intestine of *L. jeotgali* colonized *C. elegans* N2, with scale bars = 2 μm. (G) Quantification of the intestinal brush border depth from TEM images of uncolonized (N=4, n=56), *Ca*. L. limosiae colonized animals (N=3, n=35), and *L. jeotgali* colonized (N=3, n=36). (H). Quantification of the area of the intestinal lumen for uncolonized (N=4, n=20), *Ca*. L. limosiae colonized (N=3, n=17) and *L. jeotgali* colonized (N=3, n=15), animals. Statistical analyses performed by ordinary one-way ANOVA where ns = not significant and p < 0.0001 = ****.

*Ca*. L. limosiae colonization caused a significant reduction of the intestinal brush border depth, with a depth of 0.43 μm compared to the 1.026 μm depth seen in uncolonized animals (Fig. 4D-E, quantified in Fig. 4G). Effacement of the intestinal microvilli and loss of the glycocalyx are expected to impact the intestinal integrity and result in the increased intestinal leakiness seen in the “Smurf” assay (see Fig. 3I-J). By contrast, *L. jeotgali* (LUAb3) colonized animals did not display a significant change in intestinal brush border depth, with a depth of 0.98 μm (Fig. 4F and 4G). Separately, we found that both adhering bacteria caused varying levels of intestinal dilation, where the natural elliptical shape of the lumen became more circular (Fig. 4D-F, 4H). This effect was quantified by measuring the area of the intestinal lumen. We found that the intestinal dilation in animals colonized with *Ca*. L. limosiae is nearly 4-fold higher than in uncolonized animals, with a concomitant decrease in the volume of the intestinal cell. *L. jeotgali* colonized animals also had a more circular lumen, with a small but significant increase in the luminal area (Fig. 4E). This partial distension may be attributed to the presence of intact bacteria colonizing the lumen.

### Adhering microbiome bacteria replicate in the gut to colonize the entire lumen

A hallmark of gut microbiome bacteria is the ability to survive and proliferate in the host intestine. We developed an assay to visually assess the proliferation of bacteria in *C. elegans*, where axenic animals were pulse exposed to adhering bacteria prior to propagation on *E. coli* OP50 lawns and chased for colonization over the course of 48 hours. We quantified colonization by fluorescence after using 16S-specifc FISH probes for each bacterial species.^21,43^ Using this technique with *L. jeotgali* (LUAb3), we found an increase in fluorescence from 0 to 48 hours post exposure (hpe) (Fig. 5A-D). *L. jeotgali* appeared to colonize simultaneously throughout the lumen and colonized ∼99% of the population by 48 hpe (Fig. 5E). Binary fission events were observed on TEM images (Fig. 4C, 4F). This demonstrates that the bacteria are replicating in the intestines of *C. elegans* even in the absence of continuous feeding of *L. jeotgali*. Binary fission events where a bacterial cell in contact with the host glycocalyx were observed more frequently than bacteria seen dividing in the lumen (Fig. 4C, 4F). A total of 19 septation events were seen in *L. jeotgali* cells attached to the glycocalyx compared to 9 events seen in the lumen (N=3, n=36). It is likely that one attached bacterium divides latitudinally and produces a non-attached daughter cell that disperses into the lumen for subsequent adherence to the intestinal epithelia.

**Figure 5.**
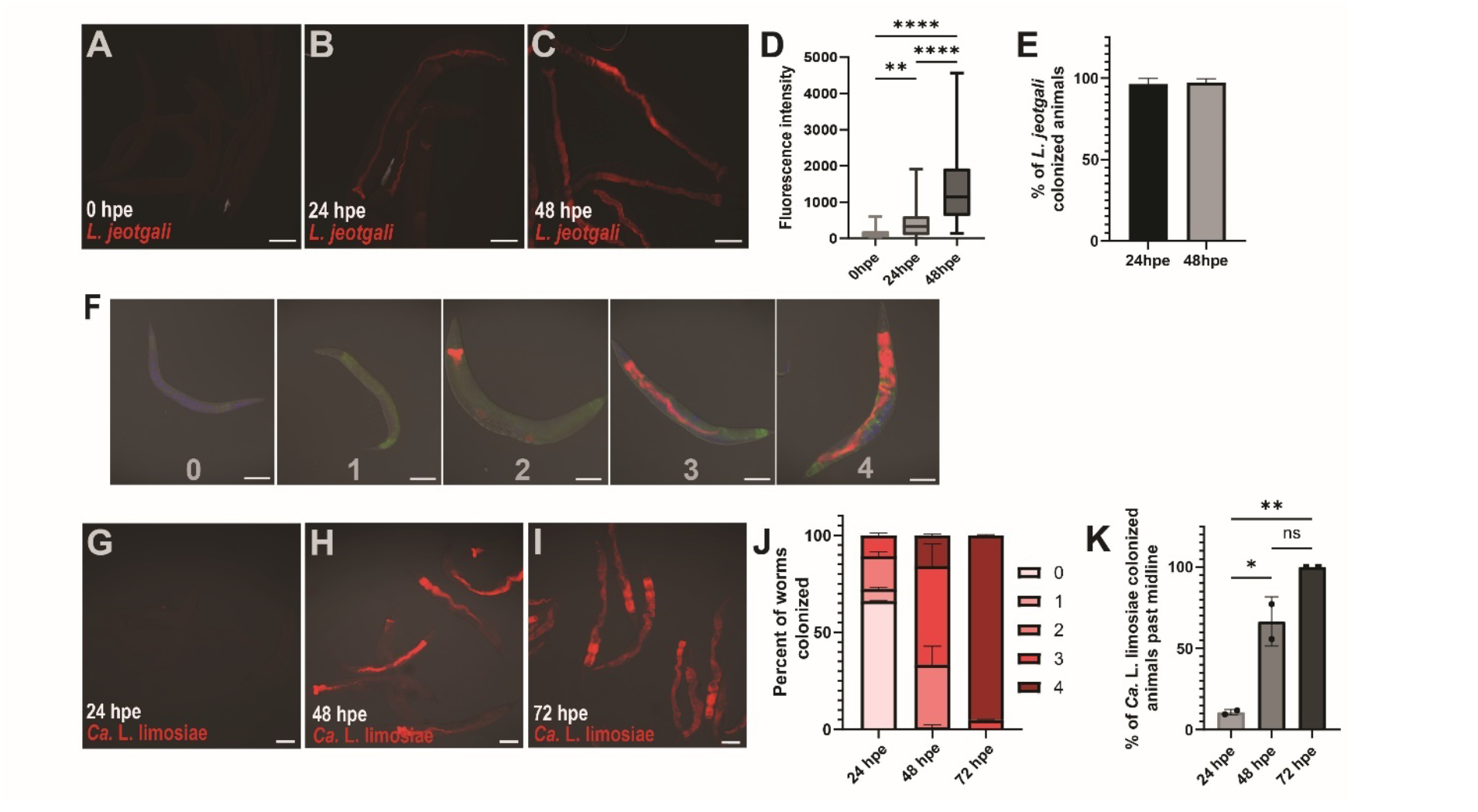
Proliferation of commensal-like *L. jeotgali* and pathogenic *Ca*. L. limosiae bacteria in *C. elegans* gut lumen. (A-C) Representative RNA FISH-stained images of *C. elegans* N2 briefly exposed to *L. jeotgali* and fixed 0 hpe, 24 hpe, and 48 hpe respectively. Red fluorescence correlates with *L. jeotgali* specific RNA FISH probes with a Cal-610 fluorophore. Scale bars are 100 μm. (D) Quantification of fluorescence intensity of *L. jeotgali* specific fluorescent probes for each fixed time point. n=30 independent in 3 independent experiments. Statistical analysis performed by multiple comparisons in a one-way ANOVA where p<0.05 = ** and p<0.001 = ****. (E) Percentage of *C. elegans* N2 colonized with *L. jeotgali* in the intestine at 24 hpe and 48 hpe, where N=3, n=30. (F) Categories of directional *Ca*. L. limosiae colonization, where 0 represents no colonization, 1 represents bacteria present only in the anterior end, 2 is bacteria detected present between the anterior and midline of the animal (vulva), 3 is bacteria found from the anterior end past the midline, and 4 is bacteria colonizing the entire anterior-posterior length of the intestine. Scale bars are 100 μm. (G-I) Representative RNA FISH-stained images of *C. elegans* N2 exposed to *Ca*. Lumenectis limosiae briefly and fixed at 0 hpe, 24 hpe, and 48 hpe. Scale bars are 100 μm. (J) Percent of *Ca*. L. limosiae colonized animals in each geographical category for each time point. (K) Quantification of the percentage of animals that were colonized past the midline (categories 3 and 4) in each time point. n is at least 90 animals in two independent experiments. Statistical analysis performed by multiple comparisons in a One-Way ANOVA where p<0.05 = *, p<0.005 = **, and ns = not significant.

In a similar pulse-chase assay, *Ca*. L. limosiae (LUAb1) was observed to colonize in a directional manner, starting at the anterior end of the intestine and moving posteriorly. We quantified this progression into different categories (from 0-4) depending on the extent of anterior-to-posterior colonization, with 0 representing no bacterial colonization, 1 at the anterior end only, 2 between the anterior end and the vulva, 3 from the anterior end past the vulva, and 4 representing colonization of the entire gut lumen (Fig. 5F). *Ca*. L. limosiae colonization became more severe over time. Between 24 hours post exposure to 72 hpe, we saw a progression of *Ca*. L. limosiae colonization from the anterior end to past the midline of the intestine (Fig. 5G-I). And at 48 hpe, 100% of the animals were colonized past the midline (Fig. 5J-K). Unlike *L. jeotgali*, we observed fewer binary fission events of attached *Ca*. L. limosiae cells dividing by TEM (Fig. 4B, 4E). Instead, division occurred more frequently in the lumen with 29 septation events occurring in non-attached cells versus 10 events in attached cells (N=3, n=35). It is likely that *Ca*. L. limosiae replication happens primarily in the intestinal lumen and then new daughter cells attach to the epithelia.

Altogether, the contrast between the dynamics of colonization of *L. jeotgali* LUAb3 and *Ca*. L. limosiae LUAb1 supports the idea that these natural members of the nematode microbiome have distinct mechanisms for replication and persistence in the intestine.

### Effects of microbiome bacterial competition on the biogeography of gut lumen colonization

Colonization by beneficial or commensal bacteria in the intestine can aid in the defense of *C. elegans* against invading pathogens.^44,45^. In the first experimental paradigm, we tested whether pre-colonization with an adherent commensal can protect from colonization by an adherent pathogen. We exposed axenic *C. elegans* L2 larvae to *L. jeotgali* for 24 hours prior to challenging with *Ca*. L. limosiae at the L4 stage for 2 hours. Following the 2-hour exposure period, the samples were washed and grown on OP50 seeded NGM plates for 24 hours. Animals were then fixed, and RNA FISH stained using *L. jeotgali* and *Ca*. L. limosiae specific fluorescent probes (Fig. 6A, top). We observed that pre-colonization with *L. jeotgali* severely reduced *Ca*. L. limosiae colonization past the midline at 24 and 48 hpe (Fig. 6B-C, 6F). By contrast, *L. jeotgali* colonization showed no obvious change in colonization at 48 hpe when *Ca*. L. limosiae was added (Sup. Fig. 2).

**Figure 6.**
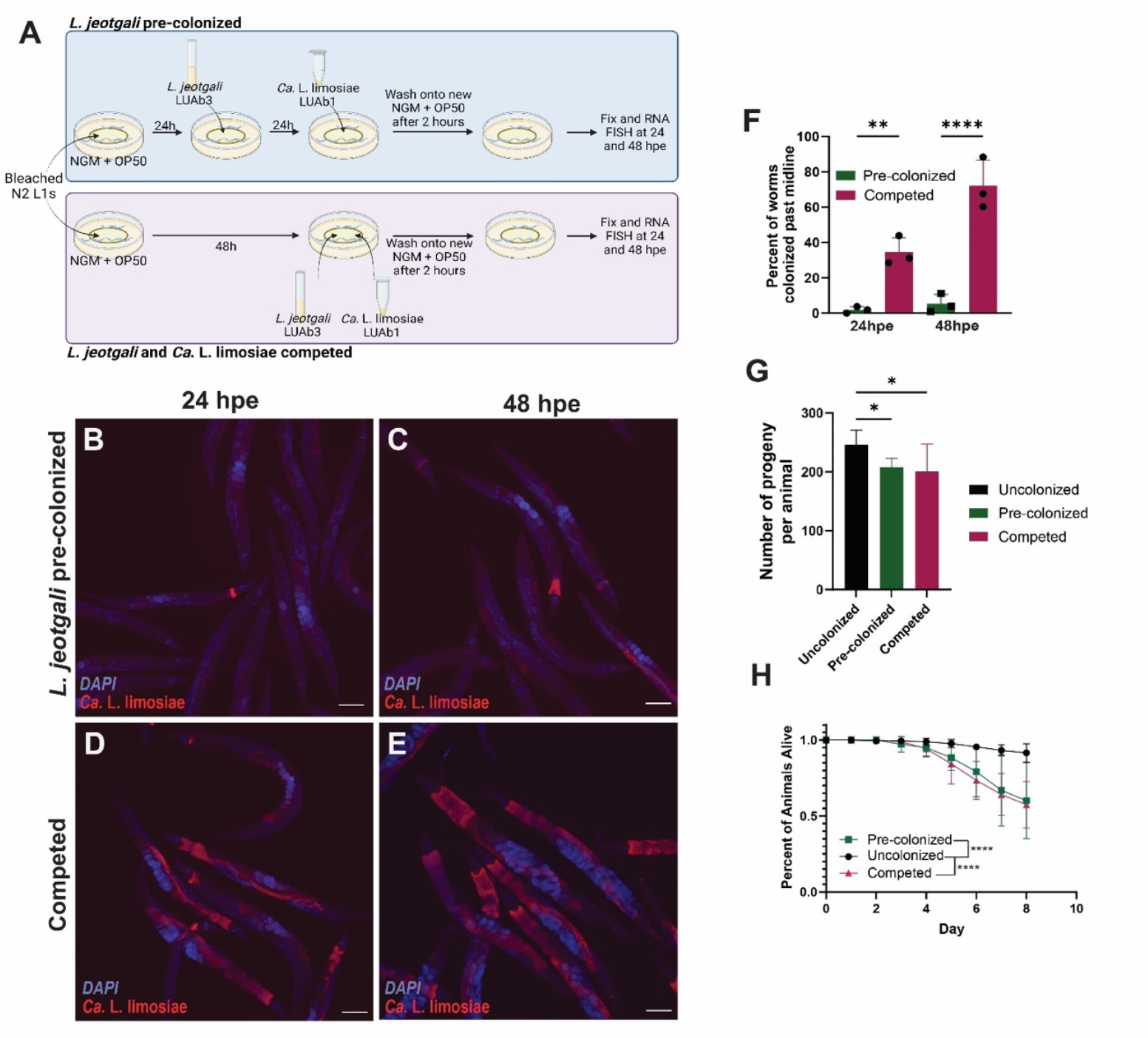
Effects of microbiome bacterial competition on the biogeography of gut lumen colonization. (A) Graphic representation of the pre-colonized and competition assay. Created with BioRender.com. (B, D) Representative RNA FISH-stained images of *C. elegans* N2 pre-colonized with *L. jeotgali* and pulse exposed to *Ca*. L. limosiae. Animals shown are fixed 24- and 48 hpe, respectively. Only the red fluorescence indicating *Ca*. L. limosiae are shown in all RNA FISH images. DAPI is shown in blue and scale bars are 100 μm. (C, E) Representative RNA FISH-stained images of N2 exposed to *L. jeotgali* and *Ca*. L. limosiae simultaneously for two hours. The animals shown are fixed at 24 hpe and 48 hpe. (F.) Quantification and comparison of the percent of *Ca*. L. limosiae colonized past the midline in either pre-colonized or competed animals at two time points. Statistical analyses performed via multiple comparisons one-way ANOVA where p<0.005 = ** and p<0.0001 = **** for three independent experiments where n is at least 98 animals. (G) Lifespan during the most reproductively active period of uncolonized, *L. jeotgali* pre-colonized, and competed animals where n=60 for 3 independent experiments. Life span curves were analyzed by Mantel-Cox test, p< 0.0001 = ****. (H) Total brood size of *L. jeotgali* and competed animals in comparison to uncolonized animals across the span of 4 days. n=60 for 3 independent experiments.

In the second experimental paradigm, we tested whether competing both adherent commensal and pathogenic bacteria simultaneously can affect pathogenic colonization. Axenic *C. elegans* L4 larvae were exposed for 2 hours to *Ca*. L. limosiae and *L. jeotgali* (Fig. 6A, bottom). Under this paradigm, we found that neither bacterium was dramatically affected in its colonization capacity by the other. At 48 hpe, ∼80% of animals were colonized by *Ca*. L. limosiae past the midline (Fig. 6D-F). At both 24- and 48 hours post exposure, we see that *Ca*. L. limosiae continues to initiate colonization at the anterior end of the intestine (Fig. 6B-E). Comparing the two experiments, *L. jeotgali* pre-colonization led to a nearly 7-fold reduction at 48 hpe in the prevalence of *Ca*. L. limosiae colonizing past the midline when compared to the simultaneous competition paradigm (Fig. 6F). These data suggest that pre-colonization with an adherent bacteria can prevent the colonization by a subsequent species.

Using these two paradigms, we next tested the effect on host reproductive lifespan and brood size. Due to the reduction of *Ca*. L. limosiae colonization past the midline seen in the pre-colonization paradigm, we predicted that the animals would fare better in reproductive lifespan and brood size assays than the animals in the bacterial competition paradigm. Surprisingly, reproductive fitness decreased similarly in comparison to uncolonized animals in both experimental situations (Fig. 6G-H). Overall, even though *L. jeotgali* pre-colonization led to a severe decrease in pathogenic *Ca*. L. limosiae colonization compared to simultaneous competition, there was no concomitant benefit to host fitness. These data suggest that either the pathogenic effects of *Ca*. L. limosiae are mitigated by the presence of commensal *L. jeotgali*, or that *L. jeotgali* has cryptic virulence that emerges when a pathogenic bacterium is present.

## Discussion

We have discovered three bacterial species that colonize the *C. elegans* gut by specifically adhering to the apical side of intestinal cells. Two of these species have commensal features as they show no significant effect on host fitness, while one of them displays clear pathogenicity. Though other animal models are used to study the gut microbiome, *C. elegans* offers a distinct advantage in its transparent body, which allows for easy visualization of the dynamics of bacterial colonization in the intestine of a whole animal. We were able to observe the organization and spatial distribution, or biogeography, of adhering bacteria and observe differences among them.

### Resistance to the gut luminal flow and the bacterial niche

Attachment to the gut cells is very efficient for colonization of the lumen. Indeed, an initial two-hour exposure window is sufficient for *L. jeotgali* and *Ca*. L. limosiae persistence and proliferation throughout the host reproductive life span. While planktonic gut microbes need to withstand the defecation cycle that may empty most of the intestinal lumen every 45 seconds, adherence is a mechanism enabling persistence in the intestine.^46,47^

Most bacteria adhere by utilizing mucus-binding proteins and/or cell surface appendices, like fimbriae or pili, that bind to the mucosal layer protecting the intestinal epithelium.^48^ Although the intestinal mucus layer in *C. elegans* is not well characterized, the presence of an electron dense layer covering the microvilli of the intestine can be seen via electron micrographs.^20,49^ This layer presumably corresponds to a glycocalyx composed of glycoproteins that protect the microvilli.

Similarly to planktonic bacteria, these bacteria may find food in the intestinal lumen, through the lysis of other bacteria or nutrients extracted from the host, as suggested by the strong reduction in host intestinal cell volume. Specific to adhering bacteria, they may feed off the mucus layer itself, comprised of primarily highly glycosylated mucins.^48^ Adherence to the mucus layer covering the intestine may thus provide a further resource advantage over less adherent species. In fact, pathogens like *Pseudomonas aeruginosa* have been shown to use mucin-derived monosaccharides as a nutrition source as the bacteria colonize the intestine in *C. elegans*.^50^ The robust persistence of our adhering bacteria in the intestine throughout the lifespan of *C. elegans* may be attributed to their ability to better persist in the mucus layer and then potentially utilizing mucins or intestinal cells for nutrition.^51^

Additionally, the host likely provides protection to adherent bacteria, as well as possibly facilitating bacterial dissemination as wild nematodes travel to subsequent rotting substrates for food. In fact, the procedure we used to clean the wild isolates involved forcing animals into dauer and harsh detergent cleaning, showing that these bacteria not only can colonize dauers but also survive within them.^28^ This suggests a possible mechanism for co-evolving a relationship with a host for dissemination, as overlapping ecologies can lead to the enrichment of certain microbes in dispersing animals, allowing for the evolution of more specific associations with a host.^52^

### Evolution of attachment in different bacterial genera

We found three distinct bacterial lineages that evolved the ability to adhere to *C. elegans* intestinal cells. The most prevalent is *Lelliottia* (Table 1), which we found in several *Caenorhabditis* strains around the world. We provide evidence that *L. jeotgali* LUAb3 displays commensal-like behavior in wild *C. elegans* and is beneficial under some conditions. First, it protects against *Ca*. L. limosiae adherence. Second, it can delay the mortal germline phenotype of some C. elegans wild isolates. Another isolate of *L. jeotgali* (JUb276), isolated from *C. elegans* strain JU3224, was also shown to suppress the mortal germline of its wild cognate strain and block Orsay virus infection.^41,42^ These findings suggest that under certain environmentally-relevant contexts *L. jeotgali* can provide benefits for its host.

The second bacterial lineage we found that displayed commensal-like behavior in the tested conditions is *Ca*. Enterosymbion pterelaium. Amazingly, genome sequencing of the LUAb2 strain placed it within the clade of Rickettsiales bacteria, which were once thought to be exclusively intracellular symbionts, comprising for instance *Wolbachia, Rickettsia* and *Ca*. Midichloria; bacteria of the *Wolbachia* genus are known to infect some parasitic nematodes.^53,54^ Recently a novel Rickettsiales bacterium, *Ca*. Deianiraea vastatrix, was found to externally colonize a ciliate protist and replicate outside the host cell.^55^ This species was found to possesses a higher capability to synthesize amino acids compared to other Rickettsiales. The discovery of *Ca*. Enterosymbion pterelaium as another extracellular symbiont opens the debate on whether these species evolved from an intracellular to an extracellular obligate symbiotic association, or whether the *Rickettsiales* ancestor was an extracellular bacterium with independent lineages evolving intracellularity.

The last bacterial lineage represented by *Ca*. L. limosiae LUAb1 was found to strongly reduce host brood size and is found within Enterobacteriaceae, like *Lelliottia*. Our genomic analysis places it as an outgroup to a clade that includes among others *Escherichia, Citrobacter, Lelliottia, Serratia, Erwinia* or *Pantoea*, bacteria that are particularly prevalent in the intestinal lumen of various animals but are also found in decomposing vegetal matter where *C. elegans* proliferates.^22,26,44^

Maintaining a healthy gut microbiome is essential to prevent gut related ailments. Commensal and pathogenic gut microbes capable of adherence are more persistent residents of the intestine. Uncovering how bacteria use adherence to colonize the gut will provide insight into how the host can control how they promote or resist adherence of commensal or pathogenic microbes to regulate their microbiome. *C. elegans* provides a simplified model system in which we can better study these complex host-microbe and microbe-microbe interactions. We present a collection of bacteria to further our understanding of bacterial adherence and colonization dynamics in vivo. This has the potential to translate to future work regarding identification of host and bacterial factors facilitating adherence, limitations of commensal bacterial adherence, and further characterization of the *C. elegans* glycocalyx.

## Acknowledgments

We thank Christian Braendle, Jonathan Ewbank, Nathalie Pujol, Vincent Debat, Allowen Evin and Lise Frézal for helping sample strains with adhering bacteria. This work was supported by NIH grant R35 GM146836 and NSF IOS CAREER grant 2143718 to R.J.L., and Rees-Stealy Research Fellowship to DER. MAF is supported by the Centre National de la Recherche Scientifique. Some strains were provided by the CGC, which is funded by NIH Office of Research Infrastructure Programs (P40 OD010440) and we thank WormBase. TEM Imaging was performed at the Microscopy Rennes Imaging Center (MRiC, Biosit, Rennes, France), a member of the National Infrastructure France-BioImaging supported by the French National Research Agency (ANR-10-INBS-04).

## Author Contributions

RJL and MAF conceived of the project. DER, RJL and MAF designed and analyzed the experiments and cowrote the paper, with consultation of all authors. DER, KP, SM, EM, JFL, MAF, and RJL conducted experiments. AS contributed intellectually and conceived the use of the “smurf” assay. GM and ON conceived the methods and conducted TEM. MAF provided novel reagents. RJL provided funding and mentorship.

## Methods

### Nematode and Bacterial Strains

*Ca*. Lumenectis limosiae was discovered in *C. briggsae* strain JU3205 found on a rotting banana stem from the University of Agricultural Sciences in Bangalore, India on December 23, 2016. To transfer *Ca*. L. limosiae to *C. elegans*, we co-cultured the cleaned *C. briggsae* colonized with *Ca*. L. limosiae with *C. elegans* ERT413 (*jySi21[spp-5p::GFP; Cbr-unc-119(+)] II. Caenorhabditis* strains containing *Ca*. L. limosiae were maintained on nematode growth media (NGM) plates seeded with *E. coli* OP50 for the food source at 20°C under standard conditions. Isolation of *Ca*. L. limosiae was performed via an adapted protocol for Microsporidia spore preparation.^24^ We collected and washed gravid adult *C. elegans* N2 colonized with *Ca*. L. limosiae in M9 and lysed 100 μl of worm pellets through bead beating with silicon beads. The lysate was filtered using 5 μM filters to remove lysed nematode debris, and aliquoted into 125 μl. We added 25 μl of 10% glycerol to each aliquot and stored the preps at -80ºC for further use.

LUAb2 was found in *C. tropicalis* JU1848 isolated from rotten palm tree fruits sampled in the Nouragues Forest in the French Guiana on November 22, 2009. It was transferred to *C. elegans* ERT413 using the same co-culturing method described for *Ca*. L. limosiae. Strains containing *Ca*. E. pterelaium were maintained on NGM plates with OP50 at 20°C under standard conditions. LUAb2 did not withstand the bead-beating and filtration protocol described above for *Ca*. Lumenectis limosiae.

*L. jeotgali* LUAb3 was isolated from wild *C. elegans* LUA21 which was found on a rotting leopard plant stem (*Ligularia tussilaginea*) collected on March 18, 2019 from the San Diego State University campus. After selective cleaning of LUA21 (see below), non-*E. coli* OP50 colonies grew on the nematode growth media (NGM) plates in the absence of *C. elegans*. These colonies were isolated via streak plating. *L. jeotgali* cultures were grown at 32°C in LB media prior to use for experimentation. Animals colonized with *L. jeotgali* were maintained on NGM plates at 20°C.

### Selective cleaning of wild Caenorhabditis strains

To enrich for the adhering bacteria within the intestine, while removing external contamination, we selectively cleaned the wild *Caenorhabditis* strains as described previously.^28^ In summary, animals colonized with adhering bacteria were forced into dauer to seal off the intestine. Dauers were then washed overnight (ON) in an M9 buffer solution containing 0.25% SDS, 50 μg/mL of carbenicillin, 25 μg/mL of kanamycin, 12.5 μg/mL of tetracycline, 100 μg/mL of gentamycin, 50 μg/mL of streptomycin, 37.5 μg/mL of chloramphenicol, and 200 μg/mL of cefotaxime. Once the antibiotic and SDS solution was removed, single clean dauers were propagated to individual NGM plates. Colonization with adhering bacteria was verified through DIC imaging or RNA FISH.

### Genome sequencing and assembly

Cleaned N2 with *Ca*. L. limosiae strain LUAb1 or *Ca*. E. pterelaium LUAb2 were grown on standard 10 cm NGM + OP50 plates. We harvested 10 plates in 15 ml tubes, washed 3 times with H2O and isolated total genomic DNA using Qiagen DNeasy Blood and Tissue kit. Whole genome sequencing and assembly was conducted largely as described previously.^56^ Briefly, DNA libraries were made using Illumina Nextera DNA Library and sequenced on Illumina MiSeq at 250 bp paired end reads. Reads were process with cutadapt v3.4 using the following parameters, cutadapt -u 4 -u -6 -U 4 -U -6 --no-indels -q 15,10.^57^ Then, bowtie2 v2.5.4 was used to map away *C. elegans* reads (genome WBcel123) followed by *E. coli* OP50-1 reads (ADBT01.1) with the parameters --fast --no-mixed -X 2000 --un-conc-gz.^58^ Genomes were assembled with Spades v3.15.5 using ‘careful’ parameter and testing 21, 33, 55, 77, 99, and 127 kmers.^59^ Annotation was conducted with prokka v1.14.0 with the parameters ‘addgenes’ and an evalue of 1e-06.^60^ This Whole Genome Shotgun project has been deposited at DDBJ/ENA/GenBank under the BioProject accession PRJNA116858 and BioSample SAMN44052871 for LUAb1 and SAMN44052872 for LUAb2. This Whole Genome Shotgun project has been deposited at DDBJ/ENA/GenBank under the accession JBIEOG000000000 (LUAb1) and JBIEOF000000000 (LUAb2). The version described in this paper is version JBIEOG010000000 (LUAb1) and JBIEOF010000000 (LUAb2).

*Lelliottia spp*. strains were cultured from a single colony in LB and DNA was purified using the Qiagen DNeasy Blood and Tissue Kit. Whole genome sequencing was conducted on Oxford Nanopore R10.4.1 flow cells after v14 Library Preparation by Plasmidsaurus. Reads were filtered and downsampled to 250 Mb using Filtlong v0.2.1 (default parameters) and a rough sketch of the assembly was made with Miniasm v0.3. Using information acquired from the Miniasm assembly, reads were downsampled to ∼100x coverage with heavy weight applied to removing low quality reads.^61^ Flye v2.9.1 was used for genome assembly with parameters selected for high quality ONT reads.^62^ The assembly was polished with Flye via Medaka v1.8.0 using the reads generated from the sketch assembly. Annotation was conducted using Bakta v1.6.1.^63^ Genome completeness and contamination was analyzed by CheckM v1.2.2.^64^ This Whole Genome sequencing project has been deposited at DDBJ/ENA/GenBank under the BioProject accession PRJNA1168587 and BioSample accession SAMN44052892 (LUAb3), SAMN44052893 (LUAb14), SAMN44052894 (LUAb15), SAMN44052895 (LUAb16).

### Phylogenomic analysis

Phylogenomic analysis was conducted largely as described previously^56^, with the UBGC pipeline, using 92 up-to-date bacterial core genes isolated from the genomes analyzed.^65^ Genomes were chosen based on preliminary 16S phylogenetic tree analysis to see the predicted, closest related species. Alignments were made using UBCG (version 3.0) with parameter -a aa (using RAxML alignment based on amino acid sequences). Trees were made in MEGA X v10.0.5 using maximum likelihood with 500 bootstraps using AA substitution via JTT model and tree inference using Nearest-Neighbor-Interchange. *Ca*. L. limosiae strain LUAb1 and *L. jeotgali* strain LUAb3 were compared to the genomes of several divergent species in Enterobacterales. *Ca*. E. pterelaium was fairly divergent from any known species and was compared to the genomes of representative species in every Order in the Class Alphaproteobacteria.

### RNA Fluorescent in situ hybridization (FISH)

RNA FISH was performed as described previously.^21^ Animals colonized with their respective adhering bacteria were collected and washed in M9 + 0.05% TritonX. Following an additional wash in 1x PBS + 0.1% Tween-20 (PBS-T), animals were fixed for 30-45 minutes in 4% paraformaldehyde (PFA). Samples were washed four times with PBS-T to remove PFA and then put in a thermal shaker at 46°C at 1200 rpm overnight in hybridization buffer (900 mM NaCl, 20 mM Tris pH 7.5, 0.01% SDS) containing species-specific and universal bacterial FISH probes. Following overnight hybridization, animals were pelleted down and washed with wash buffer (hybridization buffer + 5 mM EDTA) once. Samples were then pelleted down and incubated in wash buffer for two 30-minute periods in a thermal shaker at 48 °C, 1200 rpm. After the second 30-minute incubation, the animals are pelleted and washed in PBS-T. Animals are mounted in Vectashield, antifade mounting media with DAPI.

The FISH probes target the 16S rRNA of bacteria and are conjugated to either 5-Carboxyfluorescein (FAM) or CAL Fluor Red 610 (CF610). Probes used are universal bacterial EUB338 (GCTGCCTCCCGTAGGAGT), *Ca*. L. limosiae specific 16S b001 (GAAAATAAGTATATTACCCTTATCTCC), *Lelliottia*-specific 16S b003 (CTCTCTGTGCTACCGCTCG), and *Ca*. E. pterelaium-specific 16S b002 (TGTACCGACCCTTAACGTTC).

### Fitness assays

Prior to the lifespan assay, bleached *C. elegans* ERT413 were grown to the L2 stage on 10 cm NGM plates seeded with OP50.^66^ For *Ca*. L. limosiae and *L. jeotgali* lifespan assays, L2 larvae were exposed to either one aliquot of *Ca*. L. limosiae lysate preps or 500 μL of an overnight of *L. jeotgali* overlaid onto the NGM + OP50 plate containing the L2 larvae. The animals were grown overnight at 20 °C until they reached the L4 stage. LUAb2 colonized animals were propagated on NGM + OP50 plates until they reached the L4 stage. Uncolonized worms were bleached and fed only OP50. Assays were performed in technical triplicate over three separate trials with n = 20 animals per 6 cm NGM plate. Animals were scored every 24 hours and were considered dead if they did not move following touch with a pick. The animals were picked onto new NGM plates every 24 hours and were maintained at 20 °C. Death due to user handling and missing animals were censored from the final count. Life span assays were conducted over the course of the reproductive period, 10 days. Statistical analysis was performed using Mantel-Cox test by Oasis 2.^67^ GraphPad Prism (version 10.1.0 (316)) was used to make the life span curves.

Brood size assays were performed similarly, except n = 20 animals were divided across four 3.5 cm NGM plates. The adults were moved to new NGM plates seeded with OP50 every 24 hours. Only the hatched, and thus viable, progeny were counted every day over the course of 4 days. Data was analyzed by unpaired two-tailed t-test using GraphPad Prism (version 10.1.0 (316)).

### Mortal germline phenotype assay

Wild *C. elegans* strain JU775 was grown on NGM plates seeded with OP50 at 15 °C. Three L4 larvae were transferred to NGM plates seeded with either OP50 or *L. jeotgali* LUAb3 and then grown for six days at 15 °C. Three adults from the new generation were transferred to a new NGM plate and then grown at 25 °C to begin the assay, either on OP50 or on *L. jeotgali* LUAb3. Three adults from each of the subsequent generations were transferred to new plates seeded with either OP50 or *L. jeotgali* LUAb3 until no larva was produced. The number of days to sterility was compared using unpaired two-tailed Wilcoxon rank tests.

### Transmission electron microscopy

High pressure freezing and transmission electron microscopy was conducted as described previously.^49^ Briefly, N2 animals and clean ERT413 animals colonized with *Ca*. L. limosiae (LUAb1) were grown under standard NGM growth on OP50. Animals were colonized with *L. jeotgali* by overlaying an ON culture of bacteria on standard NGM + OP50 plates. Samples were subjected to high-pressure freezing (HPM live μ, CryoCapCell) followed by freeze substitution, flat embedding, targeting and sectioning. Each adult animal was sectioned in five different places, every 10 μm and ultrathin sections (70 nm) were collected on Formvar-coated slot grids (FCF2010-CU, EMS). Grids were observed using a JEM-1400 transmission electron microscope (JEOL) operated at 120 kV, equipped with a Gatan Orius SC 200 camera (Gatan) and piloted by the Digital Micrograph program.

### Proliferation assay

An ON culture of *L. jeotgali* was grown in LB broth at 32 °C prior to the start of the proliferation assay. Bleached and synchronized *C. elegans* N2 L4s on NGM plates seeded with OP50 were overlaid with 200 μl of *L. jeotgali* culture with an OD = 2. After two hours of exposure to *L. jeotgali*, the L4s were harvested with M9 + 0.05% Triton-X and washed twice in M9. Once washed, the animals were plated on new 10 cm NGM plates seeded with OP50. At 24 hpe, the animals were harvested and washed again. Half of the washed animals were plated on another 10 cm NGM + OP50 plate, and the remaining half were fixed, and FISH stained using the b003 FISH probe. At 48 hpe, the animals were collected, washed, fixed, and FISH stained again. To quantify the percentage of animals colonized with *L. jeotgali*, bleached *C. elegans* N2 were prepared as described for proliferation assay. The number of animals where *L. jeotgali* was detected in the intestine via RNA FISH probe within the first 30 animals seen were counted at 24 hpe and 48 hpe.

*Ca*. L. limosiae proliferation was conducted similarly, except bleached N2 L2s were overlaid with one *Ca*. L. limosiae lysate prep. The animals were exposed for 24 hours, then washed and transferred to new 10 cm NGM plates seeded with OP50. At 48 hpe, the animals were washed and transferred to new NGM plates seeded with OP50 and maintained at 20 °C for another 24 hours. Animals were fixed and FISH stained with the b001 FISH probes at 24 hpe, 48 hpe, and 72 hpe.

Imaging was performed using the fluorescent Eclipse Ni microscope (Nikon) at 20x magnification. The exposure times were 200 ms for DAPI and 3 s for RFP. Fluorescence intensity was quantified as previously described (ref) using FIJI (Version: 2.14.0/1.54f). GraphPad Prism (version 10.1.0 (316)) was used to perform statistical analyses.

### Intestinal barrier function assay

*C. elegans* N2 were grown on NGM plates seeded with OP50 at 20 °C. Once the animals reached gravid adults, they were bleached with an alkaline hypochlorite solution.^66^ Synchronized L1s were transferred to 10 cm NGM plates seeded with OP50 at 20 °C and grown for 48 hours until they reached the L4 stage. L4s were collected using M9 and placed onto 10 cm NGM plates seeded with either OP50, *L. jeotgali*, or a OP50 + *Ca*. L. limosiae bacterial prep. They were exposed to the subsequent bacteria for a two-hour period at 20 °C. For each experimental condition, 60 worms were picked and transferred to four 6 cm NGM plates seeded with OP50 following the two-hour exposure. After 24 hours, the worms were transferred to new plates and grown for an additional 24 hours until they reached day 2 of adulthood.

The intestinal barrier function was assessed at day 2 via the “Smurf” assay.^39,40^ In a 96-well plate, 250 μL of a 5.0% wt/vol mixture of OP50 in LB + blue food dye (Spectrum FD&C Blue #1 PD110) was pipetted into each well. For every treatment, 30-40 worms were placed in the wells using a pick and maintained at 20 °C for three hours. The animals were then resuspended in the dye mixture and pipetted evenly onto 6 cm unseeded NGM plates. After drying in a flow hood with the lids ajar for 20 minutes, the adults were picked and transferred to 6 cm NGM plates seeded with OP50 where each worm was dropped into ∼10 μL of 15 mM sodium azide. We assessed the “Smurf” phenotype using a Nikon SMZ18 stereomicroscope under 20x and 40x magnification. The animals were scored + or - for “Smurf” phenotype. Smurf phenotype was determined by the presence of blue dye within the body cavity or both the body cavity and germline. Data analysis was performed using GraphPad Prism (version 10.2.1 (395)).

### Competition assay

Bleached and synchronized *C. elegans* N2 were grown on 10 cm NGM plates seeded with OP50 at 20 °C. At the L2 stage, 200 μL (OD = 2) of *L. jeotgali* grown in LB broth overnight at 32 °C was overlaid on top of the NGM plate to allow for *L. jeotgali* colonization. Once the animals reached L4 stage, *L. jeotgali* pre-colonized animals were overlaid with 150 μL prep of *Ca*. L. limosiae. For competition, L4 animals were overlaid with 200 μL of *L. jeotgali* culture (OD=2) and one 150 μL prep of *Ca*. L. limosiae. After a 2-hour exposure, the animals were collected from their respective plates and washed twice in M9. The animals were plated onto new OP50 seeded NGM plates. At 24 hours post exposure, the animals were collected and washed twice in M9. Half of the population was then transferred to new NGM OP50 seeded plates. The remaining population was FISH stained using red *Ca*. L. limosiae-specific 16S b001 probe and green *L. jeotgali*-specific 16S b003 probe. After 48 hpe, the remaining animals were collected and FISH stained using the probes described above. Colonization of *Ca*. L. limosiae was binned into 5 categories determined by the location of bacterial presence. 0 is no bacteria detected, 1 is bacteria present only in the anterior end, 2 is bacteria detected present between the anterior and midline of the intestine, 3 is bacteria found past the midline but the posterior end, and 4 is bacteria colonizing the entire anterior-posterior length of the intestine.

Lifespan and reproductive brood size was conducted as described in the fitness assays (see above) using *L. jeotgali* pre-colonized or *L. jeotgali*-Ca. L. limosiae competed *C. elegans* N2 at L4 stage. The fitness assays were performed in technical triplicate over three independent trials. Animals declared missing or killed due to user handling were censored. Data analysis was performed using GraphPad Prism (version 10.2.1 (395)).

### Taxonomic summary for *Candidatus* Lumenectis limosiae

#### Lumenectis defines a new genus

The Lumenectis genus comprises bacteria that colonize the lumen of the genus *Caenorhabditis* through adherence. This genus belongs in the Enterobacteriaceae family. The mode of infection is horizontal, and the site of infection is the host intestinal lumen. The etymology of the generic name,” lumen expanding” denotes the phenotype seen by DIC and electron microscopy whereby colonization by these bacteria leads to severe luminal expansion.

#### *Candidatus* Lumenectis limosiae defines a new species

*Candidatus* Lumenectis limosiae was initially found in wild *Caenorhabditis briggsae* strain JU3205, isolated from a rotting banana sampled from the University of Agricultural Sciences in Bangalore, India on December 23, 2016 (coordinates 13.0836, 77.57745). This bacterium is horizontally transmitted to other uninfected animals via the fecal-oral route and shows no evidence of vertical transmission. *Ca*. L. limosiae was transmitted to *C. elegans* by co-culturing. At the time of publication, this bacterium is not culturable in vitro. The bacterial genome was assembled in silico and phylogenomic analysis newly describe this as a novel bacterium within the new Lumenectis genus, named after the distended lumen characteristic this bacterium exhibits during colonization. Species etymology comes from the Greek goddess of starvation, Limos, as the bacterium causes a pathogenic effect on the animal. We discovered this bacterium via DIC light microscopy as bacilli attaching to the apical side of intestines. Animals colonized with this bacterium experienced symptoms of infection such as severely distended anterior lumens, sluggish movement, and clear intestinal cells. RNA fluorescence in situ hybridization (FISH) images using specific 16S rRNA probes revealed that over time this bacterium colonizes the entire anterior to posterior length of the intestine.

Although this bacterium cannot grow outside of the host, we could enrich its presence within the intestinal lumen through selective cleaning overnight. Bacterial preparations can be made by allowing the bacterium to colonize a large population of *C. elegans*, lysing the colonized animals, and removing lysed nematode debris via 0.22 um filter. These *Ca*. L. limosiae preparations could then be stored in the - 80°C for subsequent infections.

### Taxonomic summary for *Candidatus* Enterosymbion pterelaium

#### Enterosymbionaceae defines a new family

The Enterosymbionaceae family comprises a bacterium that colonize the lumen of the genus *Caenorhabditis* through adherence. This new family belongs in the Rickettsiales order. The mode of infection is horizontal, and the site of infection is the host intestinal lumen. The etymology of the family name, “intestinal symbiont” denotes that this bacterium is an organism living in close association with the host in the intestine.

#### Enterosymbion defines a new genus

The Enterosymbion genus comprises bacteria that colonize the lumen of the genus *Caenorhabditis* through adherence. This genus belongs in the Enterosymbionaceae family. The mode of infection is horizontal and the site of infection is the host intestinal lumen. The etymology of the generic name, “intestinal symbiont” denotes that this bacterium lives in close association with the host in the intestine.

#### Enterosymbion pterelaium defines a new species

The bacterium *Candidatus* Enterosymbion pterelaium was first observed in the wild *Caenorhabditis tropicalis* JU1848 strain isolated from rotten palm tree fruits collected on November 22, 2009 next to a small river in the Nouragues Forest in French Guiana. *Ca*. E. pterelaium is horizontally transmissible with no evidence of vertical transmission. *Ca*. E. pterelaium was transferred to *C. elegans* using the same co-culturing methods as for *Ca*. L. limosiae. Currently, *Ca*. E. pterelaium is not culturable in vitro. Instead, the bacterium was enriched in *C. tropicalis* and selectively cleaned before genomic DNA isolation. *Ca*. E. pterelaium could then be transferred to *C. elegans* via co-culturing with *C. tropicalis* colonized animals. Unlike *Ca*. L. limosiae, lysate from *Ca*. E. pterelaium colonized animals does not result in re-colonization of uncolonized nematodes, emphasizing an obligate nature characteristic of other members of Rickettsiales. The bacterial genome was assembled in silico, and phylogenomic analysis newly described his bacterium as a novel species belonging to the Rickettsiales order. Based on the analysis of 92 core bacterial genes we have placed this bacterium into a novel genus named Enterosymbion for its obligate symbiotic nature within the animal intestinal lumen. Due to the hair-like appearance of this bacterium, the etymology of the species comes from the Greek King Pterelaus, who possessed a golden lock of hair that rendered him immortal.

Under DIC light microscopy, this bacterium was originally discovered as thin, hair-like bacilli that directionally adhered to the intestinal epithelium of *C. tropicalis*. RNA FISH using specific FISH probes designed to the 16S rRNA of *Ca*. E. pterelaium show that this bacterium colonizes the entire length of the animal intestine. Animals colonized with this bacterium appear to be healthy, albeit with slightly distended intestinal lumens.

## Supplementary Material

**Supplementary Figure 1.**
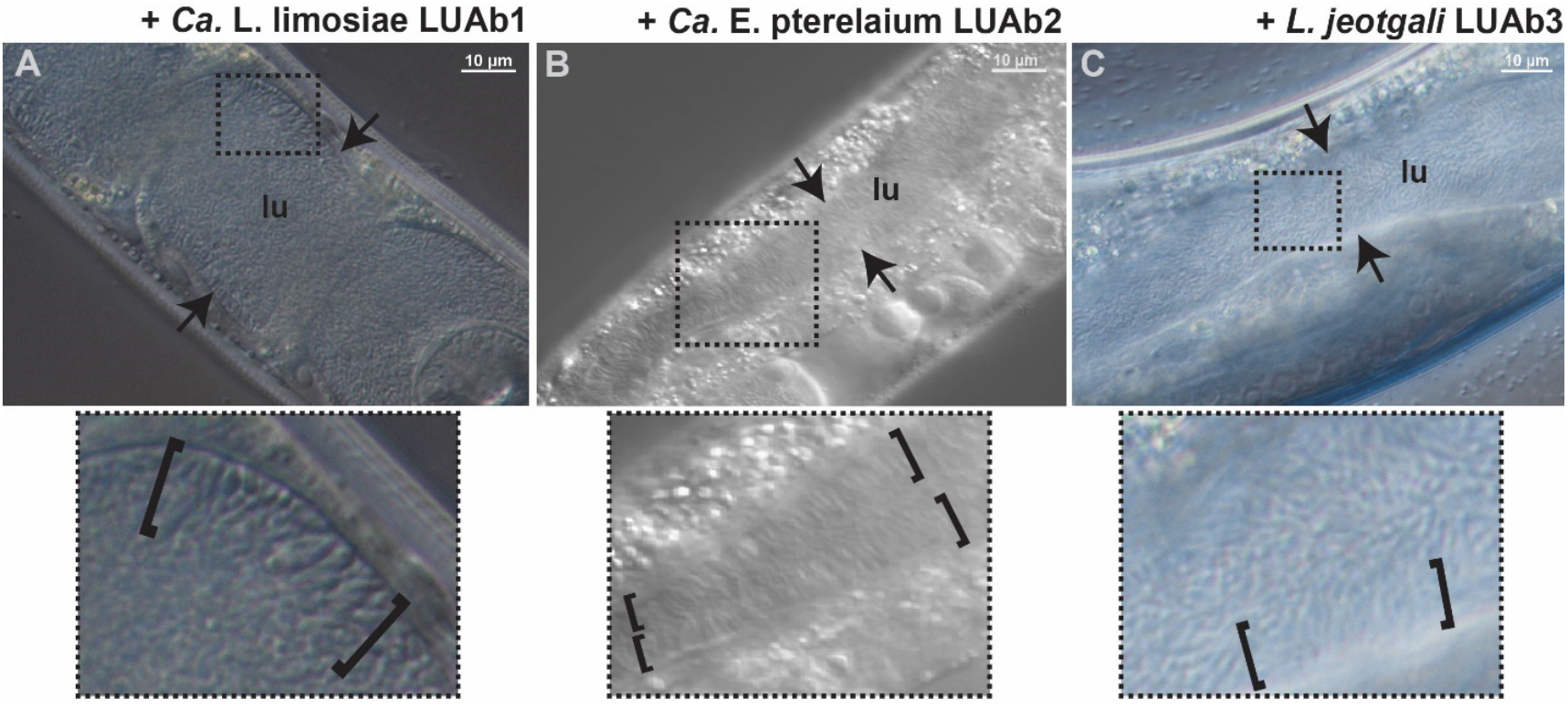
DIC images after selective enrichment of *Caenorhabditis* isolates colonized with adhering bacteria. lu denotes the lumen of the intestine, brackets indicate the adhering bacteria, and arrows denote the apical side of the intestinal epithelia. The Images below are enlarged insets of the dashed boxes above. All scale bars indicated are 10 μm. (A) *C. elegans* N2 colonized with *Ca*. L. limosiae. Note that some adhering cells appear enlarged. (B) *C. elegans* AU133 colonized with *Ca*. E. pterelaium, the thin hair-like bacilli. (C) Intestine of *L. jeotgali* colonized *C. elegans* N2.

**Supplementary Figure 2.**
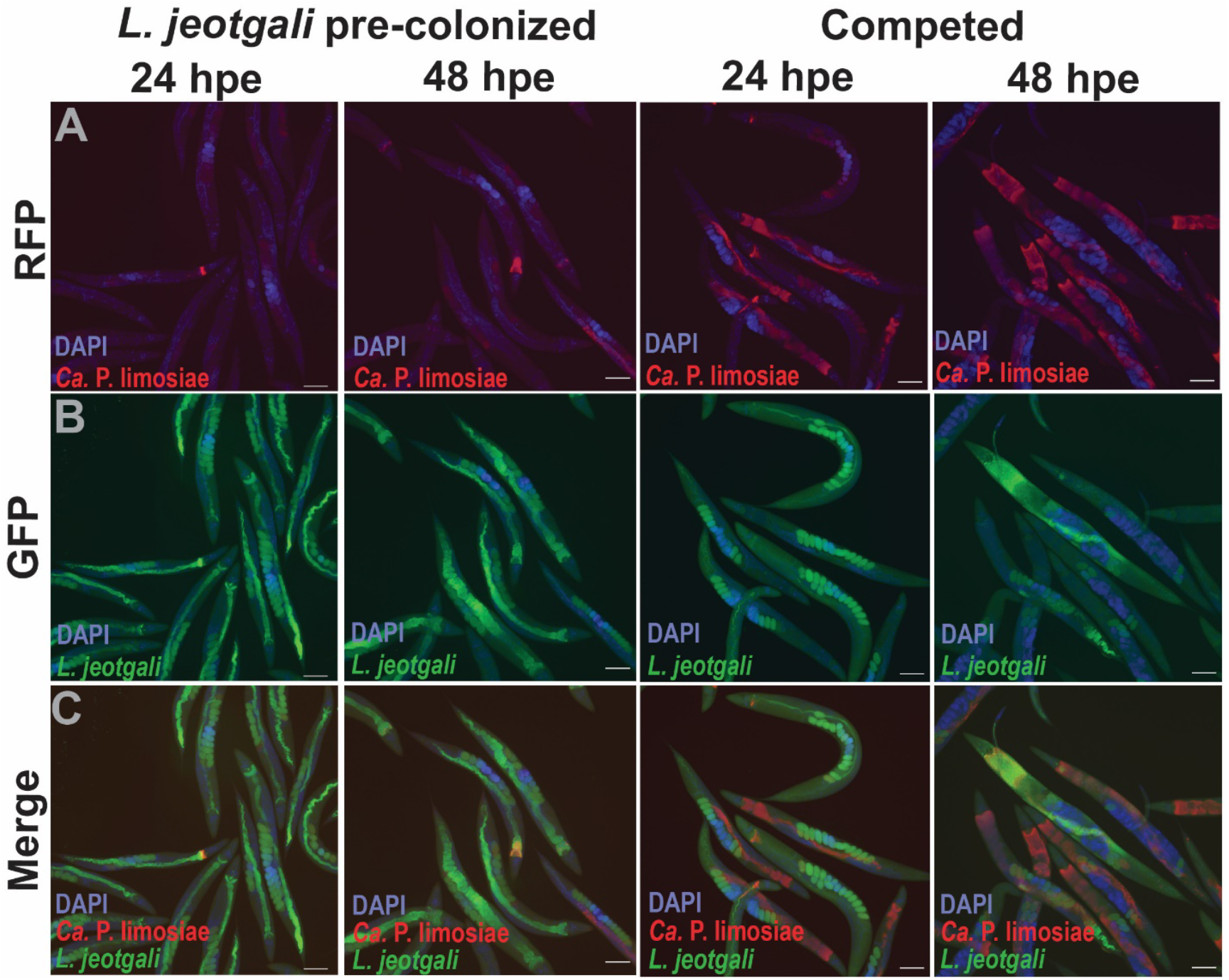
*L. jeotgali* colonization in competition assays with *Ca*. L. limosiae. Supplementary Figure 3. RNA FISH images showing the RFP, GFP, and merged channels of *L. jeotgali* pre-colonized and competed *C. elegans* N2 animals at 24 hpe and 48 hpe. All scale bars indicated are 100 μm. (A). RFP channels indicate *Ca*. L. limosiae colonization via RNA FISH as seen in Fig. 6. (B) GFP and DAPI channels indicating *L. jeotgali* colonization by RNA FISH in *L. jeotgali* pre-colonized and competed animals. (C) Merged channels of RFP, GFP, and DAPI.

